# Pooled CRISPR screens identify genes and non-coding genomic regions that regulate red blood cell density

**DOI:** 10.1101/2025.09.20.677403

**Authors:** Nicolas Brosseau, Thomas Pincez, Ken Sin Lo, Mélissa Beaudoin, Guillaume Lettre

## Abstract

Genome-wide association studies have identified >1,000 loci associated with clinically important red blood cell (RBC) traits, such as hemoglobin concentration and cell volume. However, few of these associations have been characterized at the molecular level such that most causal genes and variants remain elusive. Here, we performed pooled CRISPR screens in an erythroid cell line to identify genes and regulatory non-coding sequences that control RBC density. We perturbed 556 candidate genes and genomic sequences near 2,114 GWAS variants. We used a density gradient to detect the impact of these CRISPR perturbations on cell density. After validation, we found 17 genes and 13 regions near GWAS variants that regulate cell density. Some of these genes have previously been implicated in RBC biology (e.g. *ATP2B4*, *CCND3*, *EPOR*) although many are novel (e.g. *CHTF8*, *CTU2*, *DNASE2*). We confirmed that deletions in the osmotic stress response kinase gene *OXSR1* increase cell density, and a phosphoproteome analysis in OXSR1-depleted cells indicated that this phenotype is accompanied with a dephosphorylation of the upstream kinase WNK1 and the downstream target KCC3 (*SLC12A6*). We also combined CRISPR perturbations and RNA-sequencing to show how a non-coding genomic sequence near rs13255015 regulates the expression of the transcription factor *ZFAT* in *cis* and *SLC4A1* in *trans*. *SLC4A1* encodes Band3, a known regulator of RBC hydration and volume. Our results suggest experimental strategies to characterize GWAS findings and provide new molecular insights into the regulation of complex RBC traits.

## INTRODUCTION

Genome-wide association studies (GWAS) have identified 1000s of associations between common genetic variation and complex human diseases or clinically important quantitative traits^1^. Although these associations can be useful to develop predictive tools such as polygenic scores^2^, there are two reasons why they rarely provide molecular insights into how genetic variation influence human phenotypes. First, common genetic variants are not independent from each other because of linkage disequilibrium (LD) such that the most significantly associated variant in a GWAS locus is not necessarily the causal variant. Second, most GWAS variants map to the non-coding genome and it is often challenging to pinpoint the causal gene(s) that they regulate^3^. Many statistical and bioinformatic methods have been developed to prioritize GWAS causal variants and genes^4–13^, yet few GWAS loci have been extensively dissected at the molecular level. This molecular characterization is however essential to take advantage of the wealth of GWAS results to advance biomedical research (e.g. drug target prioritization)^14–17^.

Red blood cells (RBC) represent one of the most abundant cell-types in the human body and play a critical role in oxygen transport. The phenotypes of human RBC, such as their count, content (e.g. hemoglobin concentration) or volume vary between individuals and are highly heritable (*h*^2^∼0.25-0.64)^18^. GWAS of RBC phenotypes have identified >1,000 genetic associations, and partitioned heritability analyses of these GWAS signals suggest that the variants act cell-autonomously, likely by interfering with non-coding regulatory sequences (e.g. enhancers) active in RBC^19,20^. Because there are excellent cell culture systems to study many aspects of RBC biology and that these systems are amenable to genome editing technologies (e.g. CRSIPR/Cas9), we posited that RBC-related association results represent a unique opportunity to tackle many of the current post-GWAS challenges.

Given its fundamental role in physiology and cellular homeostasis, there are also many clinical reasons why it is important to understand the genetic factors that influence RBC traits in humans. Many human pathologies are caused or modulated by variation in RBC phenotypes. For instance, RBC hydration, which can be estimated by measuring RBC density, RBC volume (mean corpuscular volume [MCV]) or RBC hemoglobin concentration (mean corpuscular hemoglobin [MCH] and MCH concentration [MCHC]), influences two of the most common hematological disorders in the world: malaria and sickle cell disease (SCD). It is known that dehydrated RBC are less susceptible to *Plasmodium* parasite infections that causes malaria^21^. In contrast, dense RBC are associated with more severe complications in SCD patients because the polymerization of the sickle hemoglobin (HbS) depends on its intracellular concentration, which is higher in denser (dehydrated) RBC^22^. The study of rare mendelian diseases have implicated genes related to ion transport and the cytoskeleton in the regulation of RBC hydration and volume^23–25^. We have also characterized common genetic variation at a GWAS locus associated with MCHC and shown using CRISPR/Cas9 editing that it regulates an erythroid-specific enhancer for *ATP2B4*, a gene that encodes the main calcium pump in RBC^26^. Interestingly, the same *ATP2B4* genetic variants are also associated with malaria susceptibility^27^.

To functionally characterize GWAS loci associated with RBC traits and prioritize causal variants and genes associated with RBC hydration, density or volume, we performed perturbation screens in the erythroid HUDEP-2 cell line with three CRISPR/Cas9 modalities (CRISPR-KO, CRISPR inhibition [CRISPRi], CRISPR activation [CRISPRa]). Our gRNA library included 17,219 gRNA and targeted 2,114 RBC trait–associated variants and 556 candidate genes. We used a density gradient to identify perturbations that impact cell density. After a second validation screen, we confirmed 30 targets that influence HUDEP-2 density. One key finding is the identification of *OXSR1*, which encodes a kinase involved in the response to osmotic stress. We also used RNA-sequencing to profile the transcriptome of HUDEP-2 cells with CRISPR-KO edits near six GWAS variants implicated in the cell density screens to find the regulated target genes in *cis* and *trans*. Our results expand the list of validated genomic regions that carry common genetic variation that contribute to human phenotypic variation, while also providing new opportunities to identify drug targets that may be used to modulate clinically important RBC traits.

## RESULTS

### CRISPR perturbation screens for HUDEP-2 cell density

HUDEP-2 is an established erythroid cell line^28^ that we and others have used to study RBC-related phenotypes using genome editing techniques^26,29,30^. We optimized a simple centrifugation-based Percoll gradient protocol to separate HUDEP-2 based on cell density (**Supplementary Fig. 1A-B** and **Methods**). We validated this system by showing that HUDEP-2 cells that do not express the calcium ATPase gene *ATP2B4* have increased density because of increased intracellular calcium concentration, activation of the Gardos (KCNN4) channel and the resulting water loss (**Supplementary Fig. 1C**)^26^. We also confirmed that CRISPR/Cas9 edits in the coding sequence of *SLC4A1*, which encodes the cytoskeleton anchor and anion transporter protein Band3, resulted in lower HUDEP-2 cell density (**Supplementary Fig. 2**). Variants in *SLC4A1* cause different RBC pathologies in human patients, including hereditary spherocytosis^24^. While we sometimes observed variations in one of the two most extreme fractions of the Percoll gradient (#1 and #5) due to low cell numbers or cell death, the middle fractions (#2-4) robustly capture the impact of genomic perturbations on cell density (**Supplementary Fig. 1C** and **Supplementary Fig. 2**).

To identify regulators of RBC density, we designed a library of 17,219 gRNA that target GWAS variants and nearby genes associated with MCV, MCH and MCHC (**Fig. 1A**, **Supplementary Tables 1-3** and **Methods**). We performed CRISPR-KO, CRISPRi and CRISPRa perturbation screens in HUDEP-2 (**Fig. 1B**). We confirmed that gRNA targeting essential genes and introduced in the library as positive controls impaired HUDEP-2 cell proliferation in the CRISPR-KO screen (**Supplementary Fig. 3**). We also found that gRNA near 20 GWAS variants and 15 candidate genes had an impact on HUDEP-2 proliferation (**Supplementary Fig. 3B and Supplementary Table 4).**

**Figure 1.**
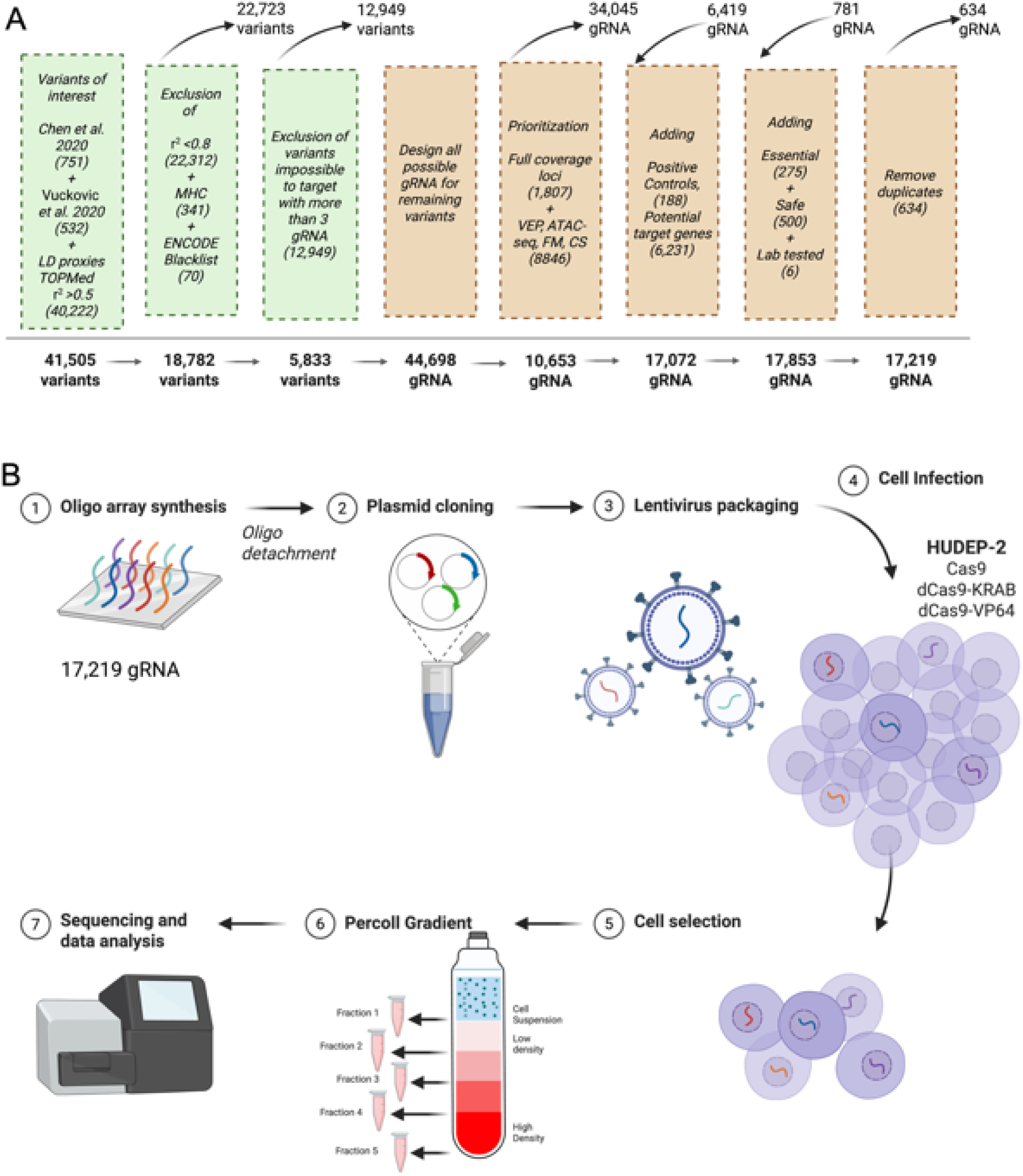
Design of the CRISPR/Cas9 perturbation screens for RBC volume. (**A**) Starting with GWAS results for RBC traits, we designed a gRNA library that includes 17,219 gRNA and that targets 2,114 GWAS variants. The different steps and filters that we applied are detailed in the **Methods**. (**B**) We used a Percoll gradient to detect the impact of CRISPR/Cas9 perturbations on the density of the erythroid cell-line HUDEP-2. We tested three Cas9 modalities: standard Cas9 which cleaves DNA and introduces insertions-deletions (CRISPR-KO), dCas9-KRAB for CRISPR inhibition (CRISPRi) and dCas9-VP64 for CRISPR activation (CRISPRa). We performed the screens using four independent biological replicates (**Methods**). MHC: major histocompatibility complex; VEP: variant effect predictor; FM: Bayesian fine-mapping; CS: chromatin state.

We analyzed the cell density data of the discovery perturbation screens by grouping the gRNA at the target level (i.e. GWAS variant or candidate gene) to maximize power and minimize the impact of gRNA off-target effects (**Methods**). Using a false discovery rate (FDR) ≤10%, we found 21, 22 and 240 targets in the CRISPR-KO, CRISPRi and CRISPRa screens, respectively (**Fig. 2A** and **Supplementary Table 4**). We grouped targets based on genomic distance (i.e. within the same GWAS locus) to determine the overlaps of the discovery screens across the three different Cas9 modalities (**Fig. 2B** and **Supplementary Table 5**). The higher number of hits in the CRISPRa compared to the CRISPR-KO and CRISPRi screens suggested a high false positive rate. For this reason, we did not characterize further the CRISPRa results.

**Figure 2.**
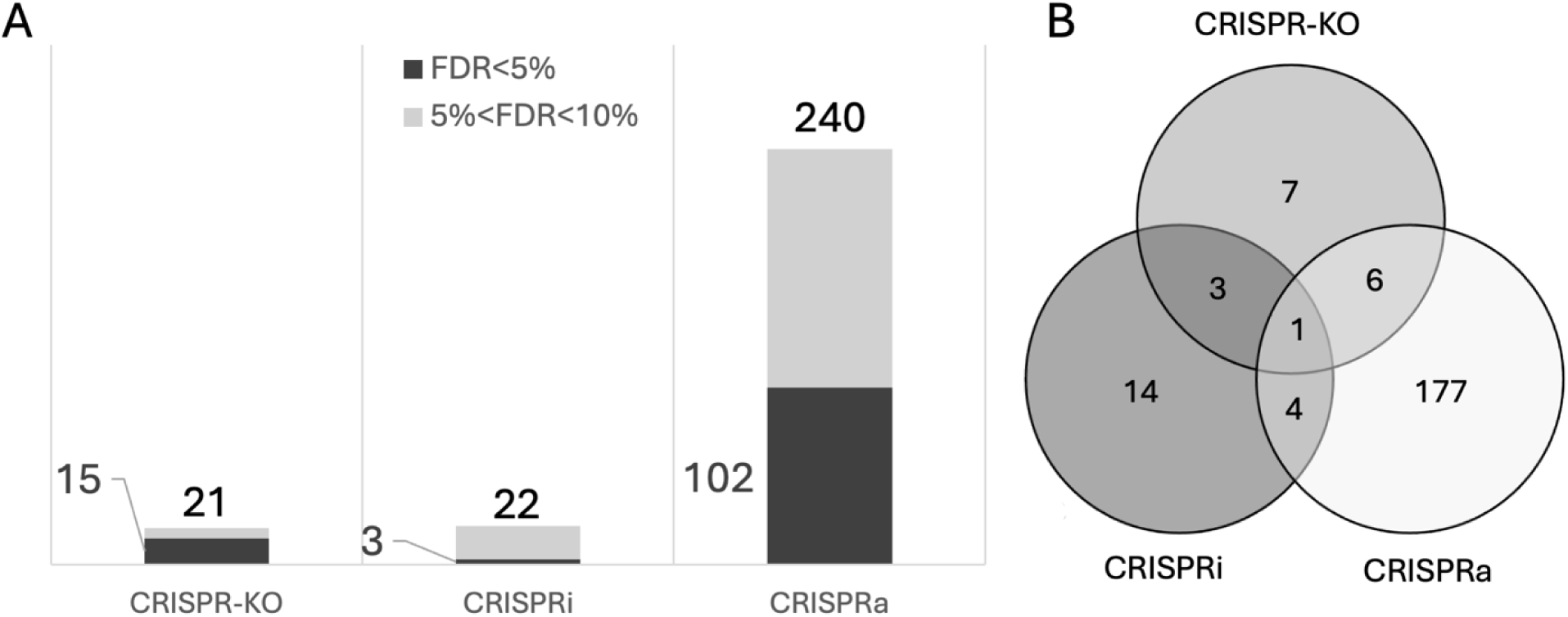
CRISPR/Cas9 perturbations identify multiple regulators of HUDEP-2 density. (**A**) Number of hits in the CRISPR/Cas9 Percoll gradient density screens with a false discovery rate (FDR) ≤5% (dark grey) or 5% < FDR ≤10% (light grey) for each Cas9 modality. The number at the top of each bar indicates the total number of hits and the number on the left of the bar corresponds to the number of hits with FDR≤5%. (**B**) Venn diagram that shows the number of genomic regions that overlap across the Cas9 modalities. For the analysis in **B**, we merged hits that target the same locus (see **Supplementary Table 5**).

### Validation screen prioritizes 30 targets that regulate HUDEP-2 cell density

To confirm some of our discovery results, we performed a validation screen using CRISPR-KO. We selected 50 targets based on several criteria: (1) statistical evidence from the discovery CRISPR-KO and CRISPRi screens, (2) overlap across Cas9 modalities, and (3) the presence of several significant targets across the same GWAS locus. We visually inspected the distributions of gRNA counts across the Percoll gradient fractions and considered the gRNA-level results to select the best 3-5 gRNA per target (**Supplementary Table 6**). To this list, we added gRNA against ten genes located near non-coding GWAS variants that were significant in the discovery screens (e.g. *TRAF4, CDH1, KLF1, MYC*). We designed and tested a library of 190 gRNA in the CRISPR-KO validation screen (**Supplementary Table 7**). We found 102 gRNA with a significant effect on HUDEP-2 cell density at FDR ≤10% **(Supplementary Table 6**). Focusing on high-confidence targets with at least two significant gRNA in the validation screen, we identified 17 genes and 13 genomic sequences located near GWAS variants, corresponding to 21 different GWAS loci, that control HUDEP-2 cell density (**Table 1**). Four (*KLF1*, *MYC*, *EPOR*, *RUVBL1*) and one (*OXSR1*) of these targets also had a negative and positive effect on HUDEP-2 cell proliferation, respectively (**Table 1**). The list of validated targets included our two positive controls (*ATP2B4*, *SLC4A1*) as well as seven other genes that have previously been implicated in erythropoiesis or RBC volume regulation (**Table 1**).

**Table 1.**
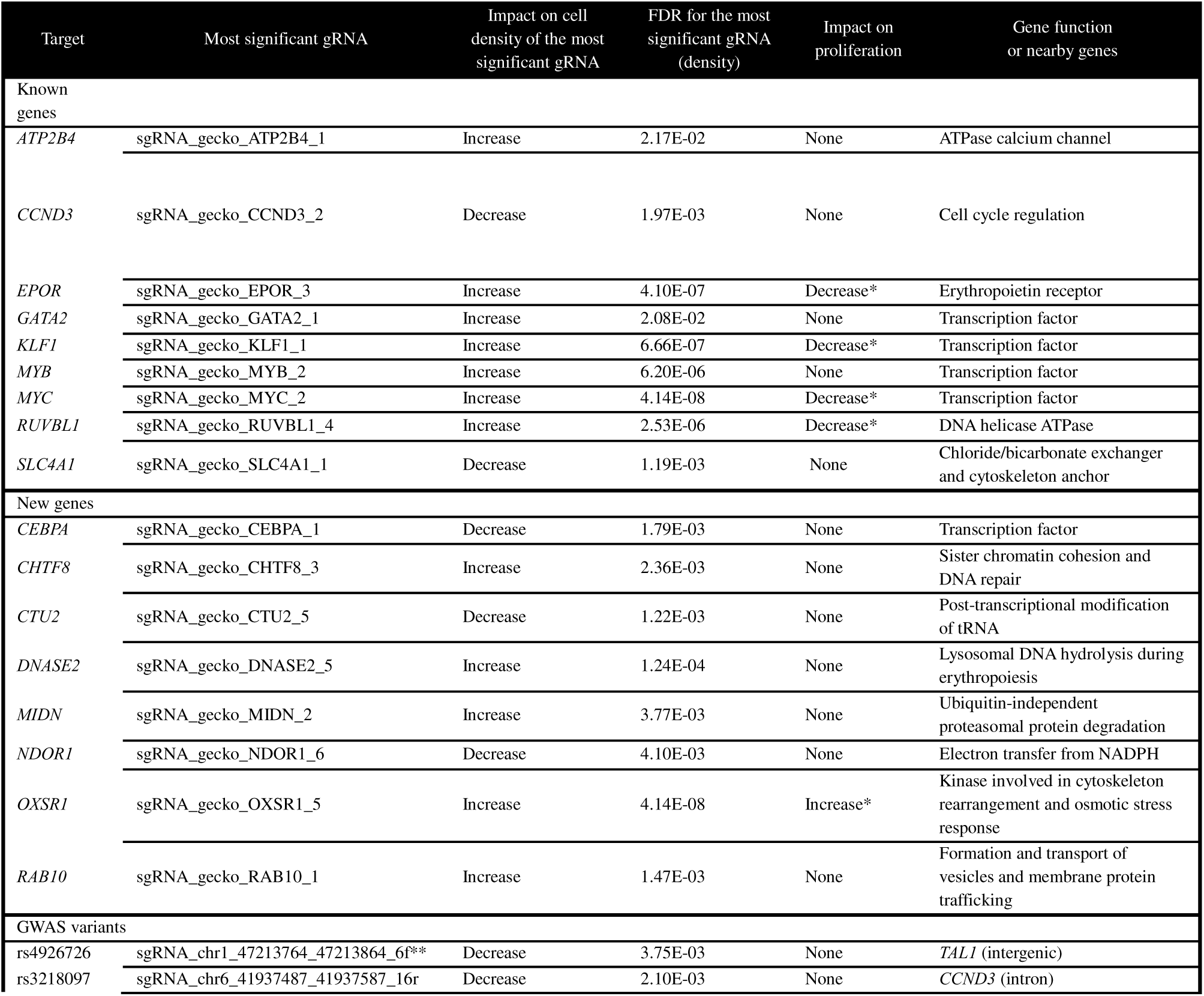
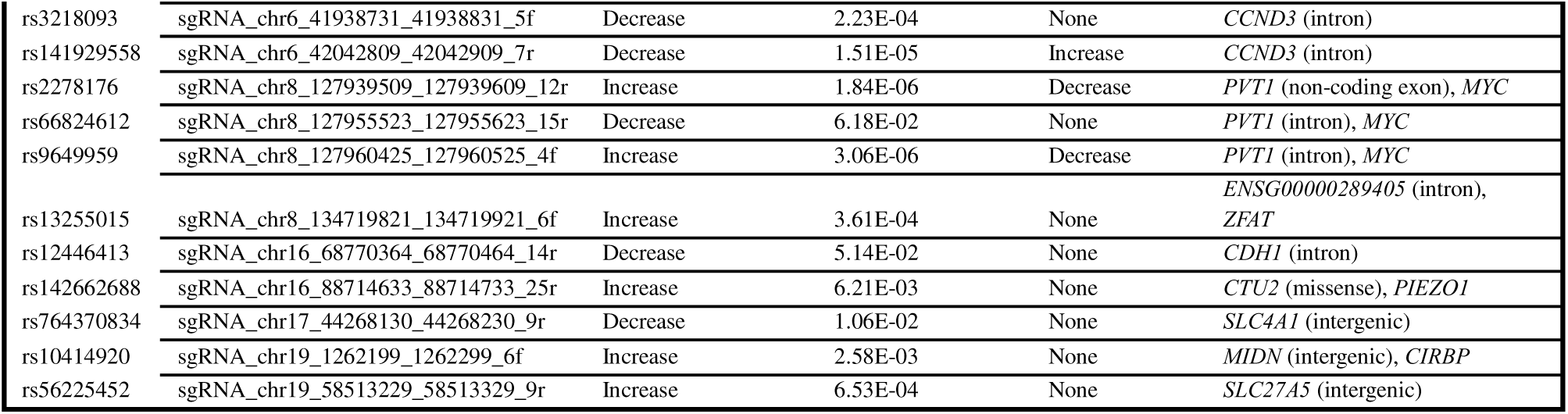
Validated targets implicated in HUDEP-2 cell density by CRISPR-KO perturbations. A list of targets (17 genes and 13 genomic regions near GWAS variants) with at least two gRNA in the validation CRISPR-KO screen (false discovery rate [FDR] ≤10%) that modulate HUDEP-2 cell density in the Percoll gradiant. All results of the validation screen are in **Supplementary Tables 4** and **6**. We defined genes as known or novel if they had previously been implicated in erythropoiesis or RBC volume regulation. *FDR<0.1 (all other gRNA that decrease or increase proliferation have a nominal p-value<0.05). **At this target, we list the second most significant gRNA because its profile in the Percoll gradient was easier to interpret to determine the direction of effect on cell density.

### *OXSR1* is a candidate causal gene at an RBC trait-associated GWAS locus

One of the strongest signals in the density screens implicated CRISPR-KO perturbations in the coding sequence of the *OXSR1* gene (**Table 1** and **Fig. 3A**). Using a single gRNA (sgRNA_gecko_OXSR1_5) in HUDEP-2 that constitutively over-express Cas9, we created a cell population with lower levels of *OXSR1* (**Fig. 3B**). We confirmed that HUDEP-2 depleted for *OXSR1* have increased cell density, validating the results from the pooled CRISPR screens (**Fig. 3C**). We included this gene in our experiment because a common indel (rs573682388) located in the *OXSR1* 5’UTR is the most likely causal RBC trait-associated variant at this GWAS locus based on Bayesian fine-mapping (**Supplementary Table 1**). However, when we targeted rs573682388 in the CRISPR screens, we did not measure a significant effect on HUDEP-2 density, suggesting that the variant is not causal or that the CRISPR perturbations at this non-coding variant were not efficient (**Supplementary Table 4**).

**Figure 3.**
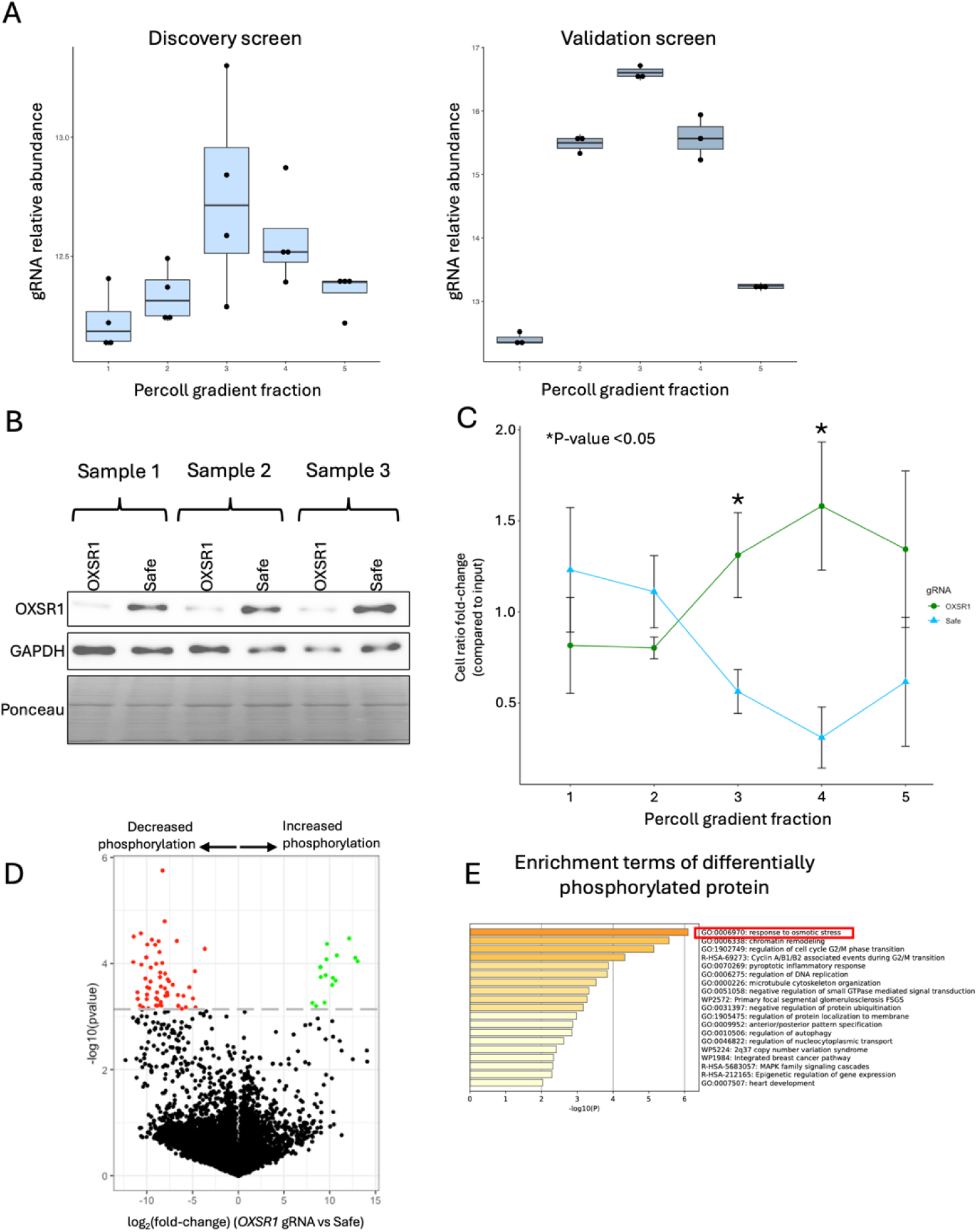
*OXSR1* deletions impact HUDEP-2 cell density and phosphoproteome. (**A**) Relative abundance of gRNA_gecko_OXSR1_5 in the different Percoll gradient fractions in the discovery (left) and validation (right) CRISPR-KO screens. The gRNA accumulates in fraction 2-5 when compared to fraction 1, indicating that *OXSR1* perturbations increase cell density. (**B**) OXSR1 immunoblotting on protein extracts from HUDEP-2-Cas9 cells that have received either an *OXSR1* or a safe gRNA. As loading controls, we present the GAPDH immunoblot results and the Ponceau staining. (**C**) Proportion of cells in each fraction of the Percoll gradient that received either the *OXSR1* targeting gRNA (green) or the safe gRNA (blue). We stained HUDEP-2 cells treated with the *OXSR1* gRNA with CellTracker Green and the cells treated with a safe gRNA with CellTracker Blue, mixed the two populations, and used flow cytometry to measure the green:blue ratio in each fraction. We used the same samples as in **B**. (**D**) Volcano plot showing differentially phosphorylated peptides in *OXSR1* KO clones as assessed by liquid chromatography and tandem mass spectrometry. The horizontal dashed line corresponds to a false discovery rate of 10%. (**E**) Enrichment terms of differentially phosphorylated proteins determined using the Metascape web tool.

*OXSR1* encodes a kinase that is part of a signalling cascade that responds to osmotic stress and regulate ion homeostasis by phosphorylating cation-coupled Cl-cotransporters^31,32^, such as NKCC1 (*SLC12A2*) in erythrocytes^33^. To gain further mechanistic insights into how *OXSR1* perturbations modulate HUDEP-2 cell density, we performed a phosphoproteome experiment. We derived HUDEP-2 clones that do not express *OXSR1* (**Supplementary Fig. 4**) and profiled phospho-peptide abundance using liquid chromatography and tandem mass-spectrometry (**Methods**). We identified 72 proteins that are differentially phosphorylated when OXSR1 is absent from HUDEP-2 (FDR ≤10%)(**Fig. 3D** and **Supplementary Table 8**). The list includes OXSR1 itself, as well as WNK1, the kinase that phosphorylates OXSR1 in response to osmotic stress^33^, and the K-Cl cotransporter KCC3 (encoded by *SLC12A6*) which is directly phosphorylated by OXSR1^34^. When we performed pathway enrichment analysis with the differentially phosphorylated proteins, we found a strong signal for proteins involved in response to osmotic stress (**Fig. 3E**). This analysis highlighted additional proteins that may regulate cell density and volume via an effect on the cytoskeleton (ABL2, KANK1).

### CRISPR-KO perturbations near 2 non-coding variants impact HUDEP-2 density and highlight candidate causal genes

The discovery and validation CRISPR-KO screens prioritized genomic regions near 13 GWAS variants in the regulation of HUDEP-2 density (**Table 1**). All but one variants are non-coding, suggesting that they regulate cell density (and RBC traits) through on impact on gene expression. We annotated these variants and found that several map to chromatin immunoprecipitation-sequencing (ChIP-seq) binding sites for important erythroid transcription factors (e.g. GATA1/2, TAL1), predicted enhancers based on chromatin states in the erythroid K562 cell line, or connected to distal genes through enhancer-promoter link predictions (**Table 2**)^35,36^.

**Table 2.**
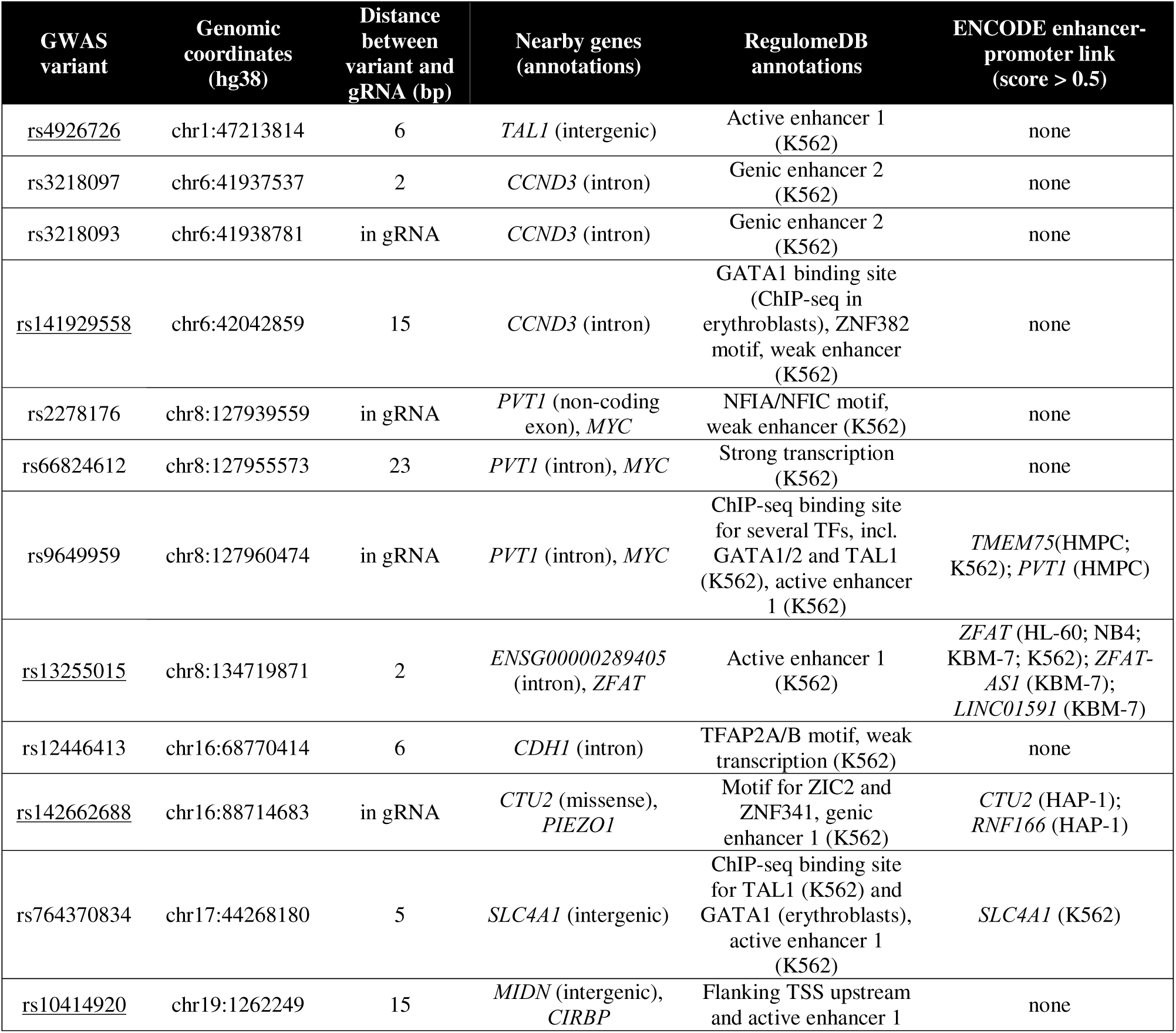
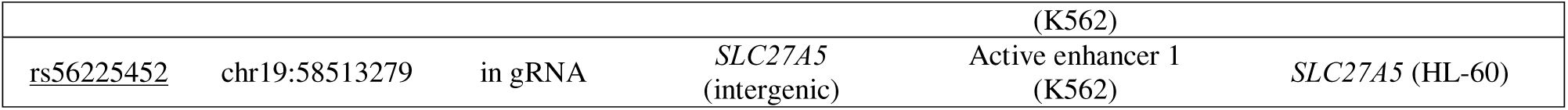
Genomic annotations of RBC trait-associated GWAS variants located near validated CRISPR-KO targets in the HUDEP-2 density screens. We annotated each variant using ENSEMBL’s Variant Effect Predictor (VEP)^54^, RegulomeDB^36^ and ENCODE’s predicted enhancer-promoter links^35^. In the first column, we underlined the 6 loci tested by RNA-sequencing. TF, transcription factor; K562, erythroleukemic cell line; HMPC, human mesenchymal precursor cells; HL-60, promyeoloblast cells; NB4, acute promyelocytic leukemia cell line; KBM-7, chronic myelogenous leukemia cell line; HAP-1, near-haploid human cell line derived from chronic myeloid leukemia.

To uncover which gene(s) these variants and surrounding genomic regions may regulate, we selected six of them and performed CRISPR-KO followed by bulk RNA-sequencing (**Supplementary Fig. S5A-G**). We chose these regions because of their validated effect on HUDEP-2 cell density in the absence of an impact on cell proliferation. For each of the target/variant, we compared the expression of genes in *cis* to the gRNA (<500-kb) between HUDEP-2 cells that have received the target gRNA vs. a safe gRNA (**Supplementary Fig. 5H-M** and **Supplementary Table 9**). For two of these regions, defined by the GWAS variants rs13255015 and rs1041920, we found genes located near the locus (<500-kb) that are dysregulated by the CRISPR-KO perturbations (**Fig. 4A-B**). For the remaining four loci, although the effect on cell density was validated and the CRISPR-KO edits detected (**Fig. S5G**), we did not identify differentially expressed genes in *cis*. As we noted before^37^, one possible explanation for this negative result may be that our bulk RNA-sequencing experiment did not have sufficient power to detect a modest effect on expression levels.

**Figure 4.**
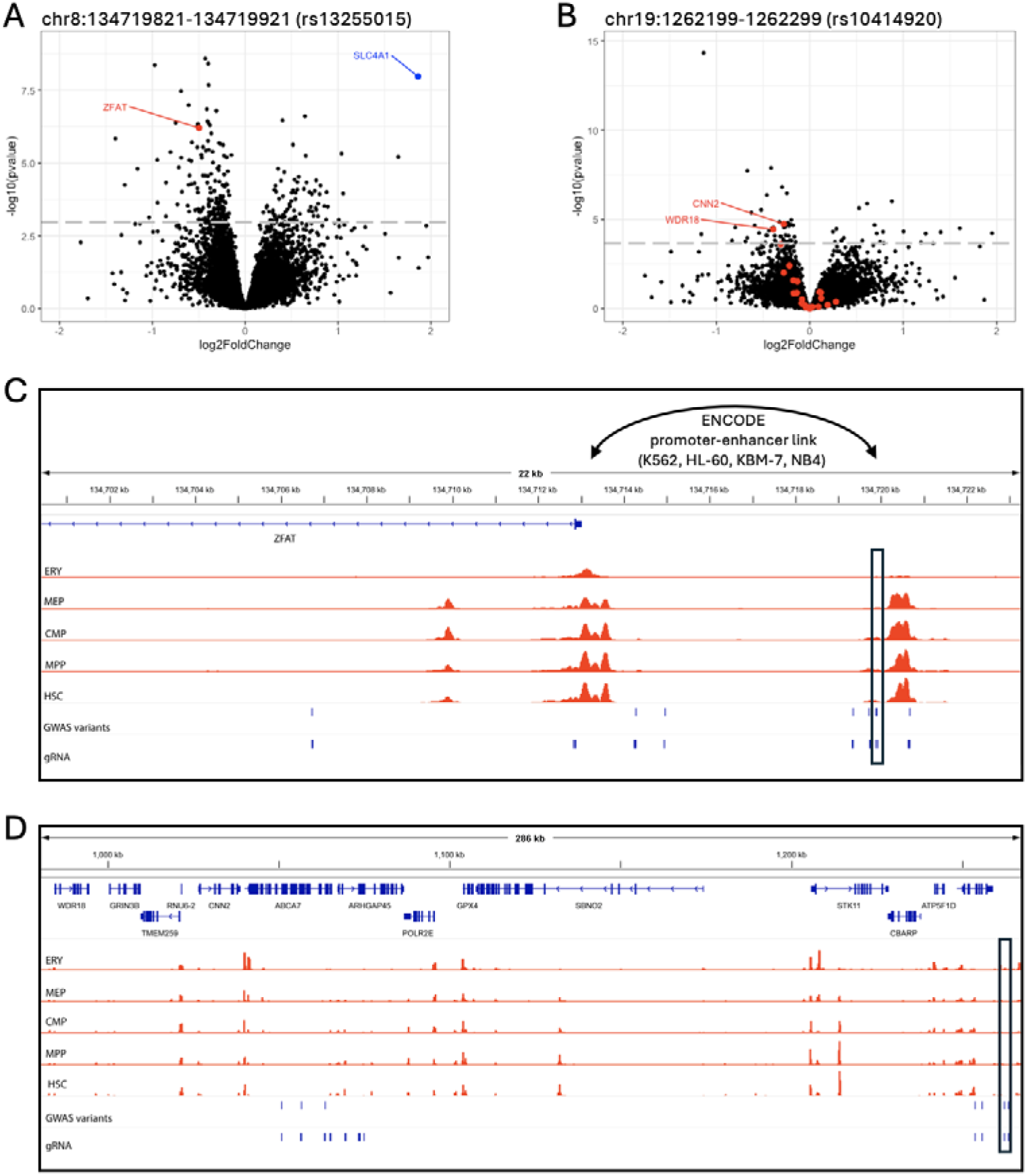
CRISPR-KO perturbations of sequences near non-coding red blood cell (RBC) trait GWAS variants impact gene expression. (**A**) Volcano plot showing differentially expressed genes in HUDEP-2 cells that received a gRNA targeting the genomic region near rs13255015. *ZFAT* (log_2_(fold-change)=-0.5, adjusted *P*=1×10^-4^) and *SLC4A1* (log_2_(fold-change)=1.8, adjusted *P*=3×10^-5^) are differentially expressed. (**B**) Volcano plot showing differentially expressed genes in HUDEP-2 cells that received a gRNA targeting the genomic region near rs10414920. *WDR18* (log_2_(fold-change)=-0.39, adjusted *P*=0.018) and *CNN2* (log_2_(fold-change)=-0.28, adjusted *P*=0.015) are differentially expressed. For **A** and **B**, genes in red are located <500-kb from the tested gRNA, and the horizontal grey line corresponds to a false discovery rate (FDR) <5%. In **A**, we also highlight in blue *SLC4A1* (Band3). (**C**) rs13255015 maps to open chromatin peaks detected by ATAC-sequencing (red tracks) in RBC precursors and is predicted to physically interact with the *ZFAT* promoter (black arc), located 7-kb downstream. (**D**) rs10414920 maps to a weak open chromatin site found in erythroblasts by ATAC-sequencing (ERY, top red track). rs10414920 is located >200-kb upstream of *WDR18* and *CNN2*. In **C** and **D**, the black box highlights the genomic region that includes the tested gRNA. ERY: Erythroid; MEP: Megakaryocyte erythroid progenitor; CMP: Common myeloid progenitor; MPP: Multipotent progenitor; HSC: Hematopoietic stem cell.

CRISPR-KO perturbations with the gRNA that targets the region near rs13255015 decreased the expression of *ZFAT*, located 7-kb downstream (**Fig. 4C**). *ZFAT* encodes a transcription factor important for erythropoiesis^38^. rs13255015 maps near an open chromatin region identified by ATAC-sequencing in RBC precursors^39^, and to an enhancer that is predicted to physically interact with the *ZFAT* promoter in several cell lines, including K562 (**Fig. 4C**)^35^. Interestingly, the gene that is the most up-regulated in this CRISPR-KO experiment is *SLC12A1* (Band3), a known regulator of RBC volume (**Fig. 4A**). The direction of the effect is also consistent: down-regulation of *SLC4A1* decreases density (**Supplementary Fig. S2**), whereas its up-regulation (via *ZFAT* down-regulation) increases cell density (**Fig. 4A**). These experiments provide a plausible molecular explanation for this GWAS signal: rs13255015 modulates the activity of an enhancer that controls the expression of *ZFAT* in *cis*, which regulates directly or indirectly in *trans* the expression of the effector RBC volume gene *SLC4A1*. For the second locus with differentially expressed in genes, the molecular mechanism is less obvious. CRISPR-KO perturbations near rs1041920 reduce the expression of *CNN2* and *WDR18*, located >200-kb downstream (**Fig. 4D**). While this genomic sequence is annotated as an enhancer in K562 and corresponds to a weak ATAC-sequencing peak in erythroid cells (**Table 2**), no links are predicted to connect it with the promoters of *CNN2* and *WDR18* (**Fig. 4D**). These two genes have also not been previously involved in RBC biology.

## DISCUSSION

In this study, we optimized a Percoll gradient that we combined with CRISPR perturbations to identify regulators of RBC density. Using this strategy, we perturbed the genomic regions near 2,114 variants associated with RBC traits by GWAS as well as 556 candidate genes. After the discovery and validation pooled CRISPR screens, we found 30 targets (17 genes and 13 variants) that regulate cell density and are confirmed by at least two independent gRNA (**Table 1**). Several of these hits validate our results as they are in or near genes already known to be involved in RBC density regulation (e.g. *SLC4A1, ATP2B4*) or erythropoiesis (e.g. *KLF1*, *MYB*, *CCND3)*. Our results also include genes that have not been previously described to influence inter-individual variation in RBC traits (e.g. *CHTF8*, *CTU2*, *DNASE2*, *RAB10*).

We identified the osmotic stress response kinase OXSR1 as a key regulator of RBC density. Using a phosphoproteome approach, we found that the kinase that phosphorylates OXSR1^40^, WNK1, is dephosphorylated in OXSR1-depleted cells, suggesting a potential feedback mechanism. Furthermore, one direct target of OXSR1, the potassium-chloride co-transporter KCC3, is also dephosphorylated in OXSR1-depleted cells. KCC3 plays a critical role in regulating ion balance, which can impact RBC hydration, volume and density. Of note, we did not detect phospho-peptides for KCC1, another direct OXSR1 target, in our mass spectrometry experiment (**Supplementary Table 8**)^41^. Our discovery screen library design included gRNA against the coding and promoter sequences of *SLC12A4* (KCC1) and *SLC12A6* (KCC3). However, CRISPR perturbations at *SLC12A4* and *SLC12A6* did not result in a cell density phenotype in our assay. We can speculate that these two ion transporters share redundant functions, and it is only by disrupting the upstream activating kinase OXSR1 that we can measure a phenotype on cell density in our screens.

Combining CRISPR-KO perturbations with RNA-sequencing yielded strong molecular hypotheses for the *ZFAT* and *WDR18*/*CNN2* loci (**Fig. 4**). In particular, we showed that disrupting a predicted enhancer that harbors rs13255015 decreases *ZFAT* expression in *cis* and increases the expression of *SLC4A1* in *trans*. *SLC4A1* encodes Band3, an anion transporter that regulates RBC hydration and volume, and severe mutations in *SLC4A1* can result in abnormally round and fragile RBC^23^. In the CRISPR-KO and RNA-sequencing follow-up experiment, we also targeted a missense variant (rs142662688) in the *CTU2* gene. *CTU2* encodes an enzyme that modifies tRNA and plays a role in maintaining genome integrity^42^. This gene was prioritized because CRISPR-KO edits with gRNA targeting different exons also impacted HUDEP-2 cell density, suggesting that *CTU2* might be the causal gene at the locus (**Supplementary Table 6**). However, we did not measure a significant effect of the gRNA targeting rs142662688 on *CTU2* expression (**Supplementary Fig. 5K**). Moreover, the gene located downstream of *CTU2* is *PIEZO1*, which encodes a mechanosensitive ion channel that regulates RBC hydration and volume, and is mutated in patients with hereditary xerocytosis that is characterized by hemolytic anemia^43^. Thus, while our data confirms that CRISPR-KO perturbations at this locus regulate HUDEP-2 density, we do not have conclusive evidence that *CTU2*, *PIEZO1*, or both genes are causal.

Our study includes several limitations. First, we carried out all experiments in an immortalized cell line. While HUDEP-2 is a robust model to study human erythropoiesis, validation in other systems could further strengthen our discoveries. Second, our screens can only detect cell-autonomous effects such that GWAS signals due to other cell-types or cell-cell interactions were missed. Third, because the discovery gRNA library was complex (17,219 gRNA), we likely missed CRISPR perturbations of weak-to-modest effect on cell density. As a result, the false negative rate of our screens is likely high. Fourth, while we tried to combine different CRISPR/Cas9 modalities, we found that the CRISPRi and CRISPRa results were less reliable. Our CRISPRa results suggest a very false positive rate for this modality in our experiments. Finally, CRISPR-KO edits do not perfectly mimic naturally occurring genetic variation such that we do not claim to have identified causal GWAS variants. Future base or prime editing experiments to install specific alleles at variants located within the prioritized non-coding genomic regions implicated in cell density could address this issue (**Table 2**).

Our results illustrate the challenge to experimentally characterize the impact of common genetic variants that have a weak effect on complex human traits. Using a simple system and a cell-autonomous phenotype, our CRISPR-based approach identified many regulators of human RBC traits. However, we also missed numerous genes and non-coding regions that modulate RBC density due to a low signal-to-noise ratio and uneven CRISPR editing efficiency. Nonetheless, our findings set the stage for future experiments, including experiments with animal models and human primary RBC, to better understand the regulators of clinically relevant traits such as RBC density, hydration and volume.

## METHODS

### Cell culture

HUDEP-2 cells were cultured as previously described^26,28^: For expansion, cells were cultured in StemSpan^TM^ SFEM (Cedarlane #09650) supplemented with 1 µg/ml doxycycline (Sigma #D9891), 0.4 µg/ml dexamethasone (Sigma #D2915), 0.1 µg/ml Stem Cell Factor (Cedarlane #255-SC-050), 3 units/ml erythropoietin (Cedarlne #287-TC-500), 2 mM L-glutamine (Life technologies #25030), 200 IU penicillin, 200 µg/ml streptomycin (Wisent Biocenter #450-201-EL) and incubated at 37°C and 5% CO_2_. Cells were always kept at a density comprised between 20,000 and 1,000,000 cells/ml. HEK293FT cells were obtained from ThermoFisher (#R70007) and cultured in DMEM (Life technologies #10569010) supplemented with 10% FBS (VWR #97068-085) at 37°C 5% CO_2_ ensuring that confluence never exceeds 80%.

### Design of the gRNA libraries

We designed our gRNA libraries using a strategy that we previously published^44^. We identified sentinel GWAS variants associated with MCV, MCH and (MCHC in two large meta-analyses of GWAS results^19,20^. We retrieved variants in strong LD with the GWAS sentinel variants in European-ancestry participants from the ARIC cohort sequenced by the TOPMed project (*r*^2^ >0.5 initially, then *r*^2^>0.8)^45^. We added variants with a fine-mapping posterior inclusion probability >0.2^19,20^. Next, we removed variants in the major histocompatibility complex (MHC) and the ENCODE black list regions^46^. In a 100-bp genomic region centered on each selected variant, we designed all possible gRNA and selected the best gRNA using the CRISPR OffTarget Tool (version 2.0.3)^47^. We prioritized variants if we could design at least four high-quality gRNA, and if the variants were likely functional based on genomic annotations (missense, nonsense, frameshift indel, essential splice site) or chromatin states in the HUDEP-2 cell line (Daniel Bauer). We added gRNA that target the coding or promoter sequence of candidate genes located near the GWAS variants^7,12,48^, the coding sequence of essential genes (positive controls), and 500 safe gRNA (negative controls). The complete list of target variants and genes is in **Supplementary Tables 1-2**, and the complete list of gRNA included in the libraries is in **Supplementary Table 3**.

### Cloning of the gRNA libraries

The gRNA libraries were cloned in pHKO-09_CM_Puro and pLenti sgRNA(MS2)_Neo^49^. Briefly, the oligo pools were obtained from Agilent, then amplified by PCR. The two vectors were digested with Esp3I (ThermoFisher #FERFD0454) and gel purified. PCR products were cloned with Gibson Assembly Master Mix (NEB #E2611L) and electroporated in Endura Electrocompetent cells (VWR #60242-2). Plasmids were endotoxin-free purified from bacteria using Machery-Nagel NucleoBond Xtra kits. Library QC was performed with Illumina sequencing.

### Cloning of single gRNA

Single gRNA were cloned in pHKO-09_CM_puro using the Target Guide Sequence Cloning Protocol from the LentiCRISPRv2 and lentiGuide-Puro: lentiviral CRISPR/Cas9 and single guide RNA rev20140722. Briefly, sequence-specific oligos (**Supplementary Table 10**) were aligned and phosphorylated, then ligated in BsmBI digested pHKO-09_CM_puro. Ligation reaction was transformed in One Shot Stbl3 Chemically Competent *E. coli* (Life technologies #7373-03). Plasmids were endotoxin-free purified from bacteria using Machery-Nagel NucleoBond Xtra kits.

### Virus production

HEK293FT cells were seeded at 8 x 10^4^ cell/cm^2^. On the next day, 68 ng/cm^2^ of pMD2.G, 104 ng/cm^2^ of psPAX2, and 136 ng/cm^2^ of plasmid library were transfected in HEK293FT cells using 1,32 µl/cm^2^ of PLUS reagent (Life technologies # 11514015) and 1,2 µl/cm^2^ of Lipofectamine 2000 (Life technologies #11668-019) in Opti-MEM (Life technologies #31985070). After 4 hours of incubation at 37°C and 5% CO_2_, the medium was changed for DMEM, with 10% FBS, and 1% BSA. Viruses were collected 2 days after.

### Infection and selection

HUDEP-2 cell infection was always carried out at a cell density of 2 x 10^5^ cells/ml in presence of polybrene (Sigma #H9268). For the screens, infection was made at a multiplicity of infection of 0.3. For single guide testing, 150 µl of viral suspension was used for each 1 x10^5^ cells. Antibiotic selection was performed 24 hours after infection in the absence of any other antibiotic, excepting penicillin and streptomycin. Puromycin (Sigma #P9620) selection was performed at a concentration of 1 µg/ml for 5 days (CRISPRi) or 6 days (CRISPR-KO). Geneticin selection (Life technologies #10131027) was performed at a concentration of 1500 µg/ml for 4 days (CRISPRa).

### CellTracker staining

HBSS/HEPES buffer was prepared by supplementing HBSS (ThermoFisher #14170112) with 1 mM CaCl_2_, 0.5 mM MgCl_2_, 10 mM HEPES pH 7.3 and 0.1% BSA. Working solution was prepared by adding either 1 µM of Cell Tracker Blue CMAC Dye (Life technologies #C2110) or 1 µM of Cell Tracker Green CMFDA Dye (Life technologies #9225) to HBSS/HEPES buffer. HUDEP-2 cells were centrifuged and resuspended in working solution at a density of 1 x 10^6^ cells/ml and incubated at 37°C 5% CO_2_ for 30 to 60 min. Cells were then washed 2 times in HBSS/HEPES buffer, then resuspended in HBSS/HEPES buffer.

### Percoll gradient procedure

Solution A (3.68% BSA in water) was transformed into Solution I by adding 1 ml of HEPES buffer (0.2 M HEPES, 2.5 M NaCl, 0.09 M KCl, pH 8.0) to 19 ml of Solution A. Solution B (3.68% BSA in Percoll PLUS Cytiva [Sigma #GE17-5445-02]) was transformed into Solution II by adding 200 µl of water and 800 µl of HEPES buffer pH 7.3. Different density solutions were prepared by mixing different ratios of Solution I and Solution II. Phenol red was added to every other layer at a concentration between 16.8 µg/ml to 19.2 µg/ml. The different density solutions were layered in a 15 ml tube (2 to 2.5 ml for each solution), and cell suspension was added at the top of the density gradient. Tubes were centrifuged at 800 x g for 30 min using soft acceleration and soft deceleration. Cells were collected at each interface and mixed with PBS in 1.5 ml tubes. The tubes were centrifuged for 1 min at 800 x g. Supernatant was removed. For FACS analysis, cells were fixed for 1 hour in a 4% PFA solution. For gDNA extraction, cells were directly resuspended in PBS. The gDNA extraction was performed with DNeasy blood and tissue kit (Qiagen #69506). The gDNA concentration was determined with a Qubit 4 Fluorometer using the Qubit 1X dsDNA HS Assay Kit (Life technologies #Q33231). If required, gDNA was precipitated using ispropanol precipitation by adding 1 volume of isopropanol, 0.1 volume of GlycoBlue Coprecipitant (ThermoFisher #AM9515), and 0.2 volume of NaCl 5 M before a 15 min incubation at RT. Samples were centrifuged 15 min at 15,000 xg. The supernatant was removed, and the pellet was washed twice with ice-cold (−20°C) 80% ethanol before 1 min air dry, then resuspended in an appropriate volume Qiagen elution buffer. New gDNA concentration was assessed with Qubit.

### RNA extraction

RNA was extracted using the Qiagen RNeasy Plus Mini kit (#74136) following the manufacturer’s instructions. Quality, purity, and concentration were assessed with Bio Tek Cytation 5 imaging reader Nanodrop from Agilent and an Agilent 2100 Bioanalyzer using the High sensitivity Agilent RNA 6000 Nano Kit (Agilent #5067-1511). For the bulk RNA-sequencing experiments, we had three biological replicates per gRNA.

### Genomic DNA extraction and TIDE PCR

The Quick Extract DNA extraction solution (Mandel Scientific #EPI-QE0905T) was used on pellets of 500,000-750,000 cells. 100 µl of Quick extract were added to cell pellets and mixed by up and down. Tubes were incubated at 65°C for 6 min, then vortexed, then incubated at 95°C for 5 min, and vortexed again. Genomic DNA extracts were diluted 1:10, then 10 µl of the dilution was used in the PCR reaction using NEBNext High-fidelity 2X PCR Master mix (NEB #M0541) following the manufacturer’s instruction with an annealing temperature of 64°C and 35 cycles. Primers used for this step are listed in **Supplementary Table 10**. PCR products were purified on gel using the QIAquick gel extraction kit (Qiagen #28704). Samples were sequenced using Sanger sequencing. The percentages of indels were determined using TIDE^50^.

### PCR preparation for NGS

The gRNA present in each collected fraction was amplified by PCR using pooled NGS-Lib_Fwd primers and one CRISPRi_rev (for CRISPR-KO and CRISPRi), or one CRISPRa_rev (for CRISPRa) primer per fraction to insert fraction specific index (**Supplementary Table 10**). The PCR was performed using the NEBNext High-fidelity 2X PCR Master mix (NEB #M0541) for 25 cycles. An annealing temperature of 63°C was used. For each PCR reaction, 500-1000 ng of genomic DNA was used. The number of reactions was adjusted according to the total amount of gDNA extracted from each fraction. If the total amount of gDNA was lower than the target amount, all gDNA was used without adjusting the reaction volume. All PCR reactions were pooled together, and the PCR products were concentrated using SPRIselect Reagent kit (Beckam Coulter #B23318) at a ratio of 1X. SPRIselect reagent/PCR mix was incubated 5 min at room temperature, then tubes were placed on magnetic stand. The supernatant was removed. Beads pellet was washed twice with 80% ethanol. DNA material was eluted by incubating the beads 2 min in Qiagen’s EB buffer. The supernatant was collected, and DNA concentration was assessed using Qubit 4 Fluorometer and the Qubit 1X dsDNA HS Assay Kit. Two µg of purified material was loaded on 2% agarose gel. Bands were excised from gel and purified using QIAquick gel extraction kit (Qiagen #28704). Purified PCR product purity was assessed with Agilent 2100 Bioanalyzer and the High sensitivity DNA kit (Agilent #5067-4626), and its concentration was assessed by Qubit 4 Fluorometer and the Qubit 1X dsDNA HS Assay Kit. For sgRNA QC, the procedure was the same but using 20 ng of plasmid library as template for the PCR reaction.

### Percoll gradient CRISPR screen analyses

We used MAGeCK to count the number of occurrences of each gRNA in the two plasmid libraries and in each of the sequenced fractions (five fractions for the three types of Cas9 modalities in each of the four experiments, i.e. 60 fractions, plus two plasmid libraires^51^. We analyzed the results both at the gRNA level and at the target level (defined by the set of gRNA for a given variant or gene). At the target level, a gRNA may have been linked to several targets. Indeed, some guides (N=594) were selected several times during the construction of the library. In this case, they were only present in a single copy in the library, like the other guides. We duplicated these 594 gRNA during the analysis at the target level as a gRNA can only be attached to a single target during the computational pipeline. We used the vst function of the DESeq2 package to normalize the gRNA counts^52^. We also normalized these counts by correcting for the sequencing run and the type of plasmid to check that these parameters did not confound the results. As these adjustments had no effect, we used the uncorrected normalized count for all subsequent analyses.

To analyze the effect of gRNA on cell proliferation, we used the mle function in MAGeCK, which calculates the maximum likelihood estimation^51^. In brief, we compared the sum of the five fractions (corresponding to all gRNA present at the end of the culture) to the count in the plasmid libraries at the target level. To analyze the effect of gRNA on density, we developed two linear mixed models, one at the gRNA and the second at the target level, then performed an ANOVA.

The gRNA-level model was:

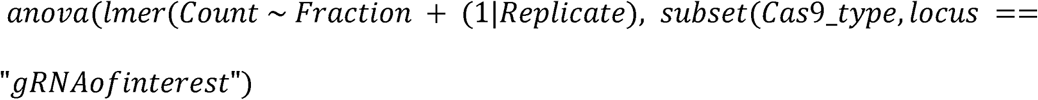

The target-level model was:

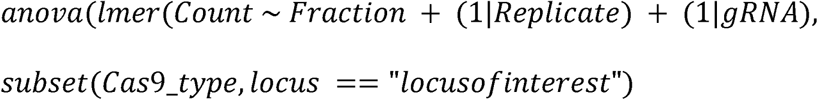

where *Count, Fraction, Replicate, gRNA,* and *Cas9type* represented the gRNA count, fraction (as a categorical variable from 1 to 5), the gRNA at target level, and the Cas9 subtype, respectively. The models aimed to identify the effect of the fraction variable for the gRNA or target of interest by adjusting for the potential effect of the experiment and the difference in the effect of the gRNA at the target level. Thus, we used a fixed effect for the *Fraction* variable and extracted the effect of this variable as our model output. We used a random effect for the variables *Replicate* and *gRNA* to obtain a better estimate of the fraction effect. We used RStudio (version 1.2.5033) and GraphPad Prism (version 9.2.0, GraphPad Software, LLC, CA) for statistical analyses. We performed only two-sided tests. We used Storey Tibshirani’s method with a false discovery rate of 5% (i.e. q-value <0.05) to correct for the multiple ANOVA test performed.

### DNA sequencing

The prepared NGS PCR products sequenced by Genome Quebec using Illumina NovaSeq 6000 SP PE 100 lanes. RNA library preparation and sequencing was performed by Genome Quebec using HMR rRNA depleted Library Preparation and Illumina NovaSeq PE100.

### Protein extraction

For every 1 million cells, 200 µl of RIPA buffer (50 mM Tris pH8, 150 mM NaCl, 0.1% SDS, 1% NP-40, 0.25% sodium deoxycholate, 1% phosphatase inhibitor cocktail 2 [Sigma #P5726], 1% phosphatase inhibitor cocktail 3 [Sigma #P0044], 1 mM PMSF, 1X protease inhibitor cocktail [Sigma #P2714]) were added to the cell pellet in a 1.5 ml tube. Tubes were put on ice and vortexed for ten seconds every minute for 5 minutes. Samples were centrifuged for 10 min at 18,000 x g at 4°C. Supernatant was collected and kept at −80°C.

### Western Blot

15 µg of proteins were migrated on an 8% acrylamide gel, then transferred to a nitrocellulose membrane (10 V, O/N). After Ponceau staining, the membrane was cut around 45 kDa to separate GADPH (36 kDa) from OXSR1 (58 kDa). Membranes were blocked in 5% milk in TBS-T (1X TBS [BioRad #1706435], 1% Tween 20 [BioRad #1610781]) for 1 hour. Membrane above 45 kDa was incubated in rabbit anti-OXSR1 (ThermoFisher #PA5-82341) at a dilution of 1:1,000 and membrane below 45 kDa was incubated in rabbit anti-GAPDH (NEB #2118S) at a dilution of 1:5,000, both in 5% milk, at 4°C overnight. Membranes were washed 3 times in TBS-T for 10 minutes, then incubated in donkey anti-rabbit HRP-conjugated (GE healthcare #PA5-82341) at a dilution of 1:10,000 at 4°C O/N. Membranes were washed 3 times in TBS-T for 10 min, then developed with SuperSignal West Pico PLUS Chemiluminescent Substrate (ThermoFisher #34580).

### Phosphoproteome analysis

Protein samples, in RIPA buffer, were shipped on dry ice at the Center for Advance Proteomics Analyses at the Institut de Recherche en Immunologie et Cancérologie (IRIC, Montreal). For each sample, 500 ug of cell lysate (measured by Bradford assay) were reconstituted in 50 mM ammonium bicarbonate with 10 mM TCEP [Tris(2-carboxyethyl)phosphine hydrochloride; Thermo Fisher Scientific], and vortexed for 1 h at 37°C. Chloroacetamide (Sigma-Aldrich) was added for alkylation to a final concentration of 55 mM. Samples were vortexed for another hour at 37°C. 10 microgram of trypsin was added, and digestion was performed for 8 h at 37°C. Samples were dried down in a speed-vac. For the TiO_2_ enrichment procedure, sample loading, washing, and elution were performed by spinning the microcolumn at 8000 rpm at 4 °C in a regular Eppendorf microcentrifuge. The spinning time and speed were adjusted as a function of the elution rate. Phosphoproteome enrichment was performed with TiO_2_ columns from GL Sciences. Digests were dissolved in 400 μL of 250 mM lactic acid (3% TFA/70% ACN) and centrifuged for 5 min at 13000 rpm, and the soluble supernatant was loaded on the TiO_2_ microcolumn previously equilibrated with 100 μL of 3% TFA/70% ACN. Flow-through samples were kept for proteome analysis. Each microcolumn was washed with 100 μL of lactic acid solution followed by 200 μL of 3% TFA/70% ACN to remove nonspecific binding peptides. Phosphopeptides were eluted with 200 μL of 1% NH_4_OH pH 10 in water and acidified with 7 μL of TFA. Eluates from TiO_2_ microcolumns were desalted using Oasis HLB cartridges by spinning at 1200 rpm at 4 °C. After conditioning with 1 mL of 100% ACN/0.1% TFA and washing with 0.1% TFA in water, the sample was loaded, washed with 0.1% TFA in water, then eluted with 1 mL of 70% ACN (0.1% TFA) prior to evaporation on a SpeedVac. The extracted peptide samples were dried down and solubilized in 5% ACN-0.2% formic acid (FA). The samples (proteomes and phosphoproteomes) were loaded on an Optimize Technologies C_4_ precolumn (0.3-mm inside diameter [i.d.] by 5 mm) connected directly to the switching valve. They were separated on a home-made reversed-phase column (150-μm i.d. by 150 mm Phenomenex Jupiter C18 stationary phase) with a 120-min gradient from 10 to 30% ACN-0.2% FA and a 600-nl/min flow rate on a EasynlC-1200 (Thermofisher Scientific, San Jose, CA) connected to an Exploris 480 (Thermo Fisher Scientific, San Jose, CA). Each full MS spectrum acquired at a resolution of 120,000 and followed by tandem-MS (MS-MS) spectra acquisition for three seconds on the most abundant multiply charged precursor ions.

Tandem-MS experiments were performed using higher-energy collisional dissociation (HCD) at a collision energy of 27%. The data were processed using PEAKS 12 (Bioinformatics Solutions, Waterloo, ON) and a human reviewed Uniprot database. Mass tolerances on precursor and fragment ions were 10 ppm and 0.01 Da, respectively. Variable selected posttranslational modifications was phosphorylation (STY).

The data were visualized with Scaffold 5.0 (protein threshold, 99%, with at least two peptides identified and a false discovery rate [FDR] of 1% for peptides). Phosphopeptide intensities were normalized with the corresponding protein intensities in the proteome. Phosphoproteome data were processed with the DEP2 package in R. Only the phosphopeptides with values in all replicates of at least one condition were kept. VSN normalization and MinProb imputation were used. We used Metascape with default parameters to perform enrichment analyses^53^.

### Bioinformatic analyses of the bulk RNA-sequencing data

An index for all cDNA was created with Kalisto 0.44.0 from the ensembl file. The alignment of the fastq files was performed with Kalisto as well. The rest of the analysis were done in R (version 4.4.2). In brief, the ENSEMBL transcript names were linked to gene name using EnsDb.Hsapiens.v86 2.99.0 and the differentially expressed gene analysis was performed with DESeq2 (version 1.46.0). Batch effect correction was applied based on cell infection dates as well as a filtering based on a minimal read count threshold of 10. Samples of cells that received the safe target gRNA were used as reference. Removal of batch effect was assessed by principal component analysis plotting. A further filtration step based on baseMean value was performed to remove low baseMean genes using a threshold of 50 to improve data visualization quality. The distance between target and gene was based on average transcription start sites of transcripts of the genes and distance of 500 kb was used as the threshold to construct the 1 Mb window around the target position. For the 5 Mb window, a distance of 2.5 Mb on each side of the target was used. The significance threshold used was an adjusted P-value of 0.05 (Benjamini-Hochberg procedure). The horizontal lines on the volcano plots were placed between the last significant gene and the first non-significant gene.

## DATA AVAILABILITY

The CRISPR screens and bulk RNA-seq data discussed in this publication have been deposited in NCBI’s Gene Expression Omnibus and are accessible through GEO Series accession number GSE308629 (https://www.ncbi.nlm.nih.gov/geo/query/acc.cgi?acc=GSE308629).

## CODE AVAILABILITY

The code used to analyze the CRISPR screen and RNA-seq data is available at: http://www.mhi-humangenetics.org/en/resources/.

## ACKNOWLEDGEMENTS

This work was funded by the Montreal Heart Institute Foundation, the Joseph C. Edwards Foundation, the Canada Research Chair Program, and the Canadian Institutes of Health Research (Project #168902) to G.L. T.P. was supported by a Charles Bruneau Foundation fellowship award and merit scholarship program for foreign students from the Ministry of Education and Higher Education of Quebec. We thank Pablo Bartolucci with help with the Percoll gradient protocol, Estelle Lecluze for supporting the bioinformatic analyses of the bulk RNA-seq data, Jennifer Zevounou for querying the ENCODE enhancer-link atlas, and Daniel Bauer for sharing chromatin state annotations for the HUDEP-2 cell line.

## AUTHOR CONTRIBUTIONS

N.B., K.S.L. and G.L. designed the gRNA discovery and validation libraries. N.B., M.B. and G.L. planned the experiments. N.B. optimized the Percoll gradient and performed all the experiments. T.P. analyzed the density gradient results. N.B. analyzed the RNAseq and phosphoproteome data. G.L. secured funding and supervised the work. N.B. and G.L. wrote the manuscript with contributions from all authors.

