## Supplementary Figures for "Pooled CRISPR screens identify genes and non-coding genomic regions that regulate red blood cell density"

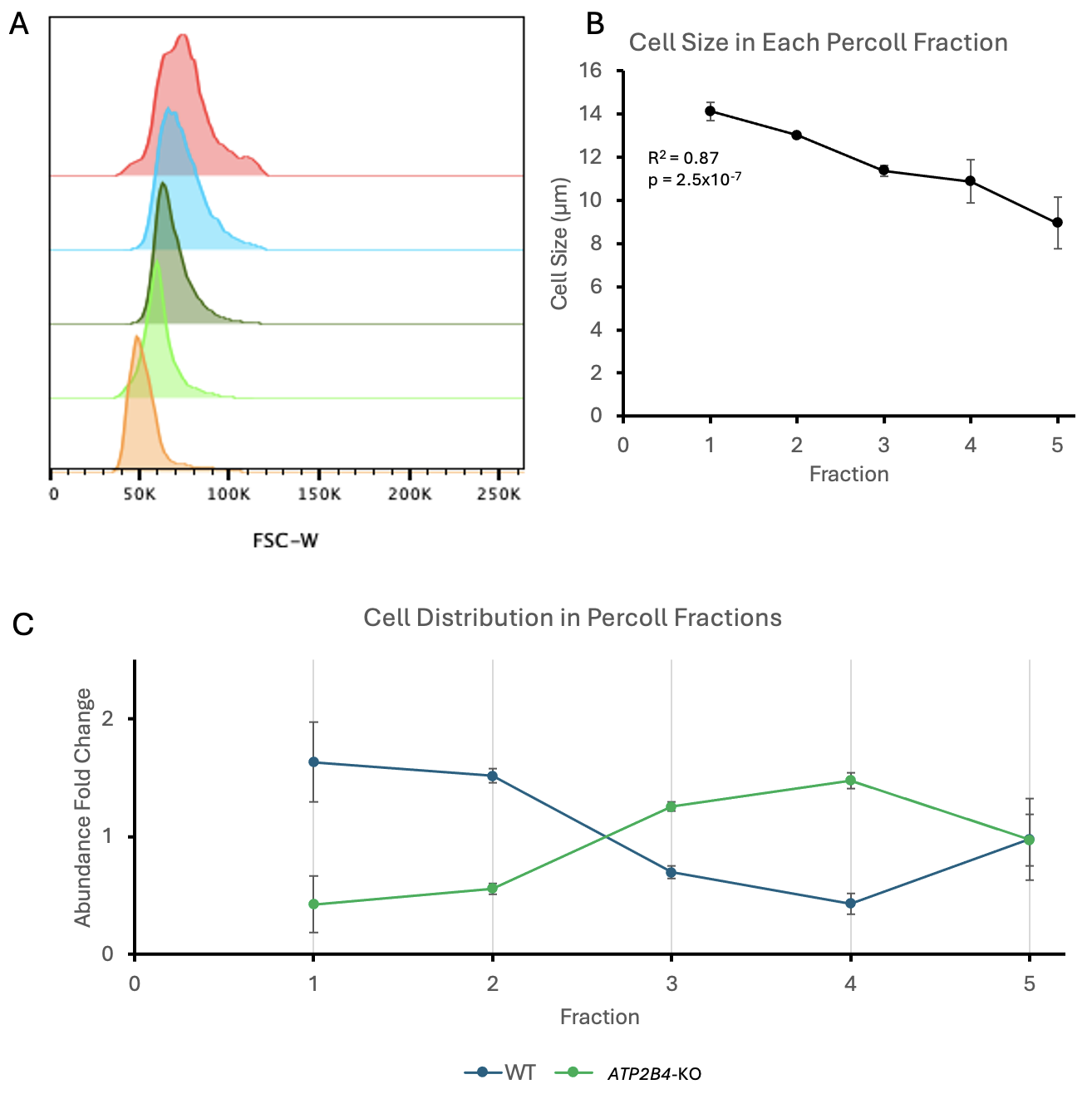

**Supplemental Figure 1. Validation of the Percoll gradient to separate HUDEP-2 cells according to their volume and density.** (**A**) FSC-W (Forward Scatter-Width) measurement of HUDEP-2 cells after separation with a Percoll gradient. After centrifugation, we collected the cells at each interface of the gradient and at the bottom of the tube. We analysed the cells by flow cytometry. Less dense, fraction #1, red; fraction #2: blue; fraction #3: dark green; fraction #4, light green; densest, fraction #5, orange. (**B**) Size of HUDEP-2 cells determined using a Countess II FL cell counter, after separation of the cells with a Percoll gradient. (**C**) Proportion of HUDEP-2 WT cells and HUDEP-2 *ATP2B4*-KO cells in each Percoll gradient fraction. We stained WT cells with a CellTracker Blue dye, and *ATP2B4*-KO cells with a CellTracker Green dye. We mixed the cells and took a sample to assess the initial WT/*ATP2B4*-KO proportion. We placed the rest of the mix at the top of a four layers Percoll gradient. After centrifugation, we collected the cells at each interface and at the bottom of the tube. We assessed the proportion of blue cells (WT) and green cells (*ATP2B4*-KO) using flow cytometry. Results are expressed as fold change compared to the sample taken before the Percoll gradient separation. The proportion of *ATP2B4*-KO cells increases in the denser fractions (from left-to-right).

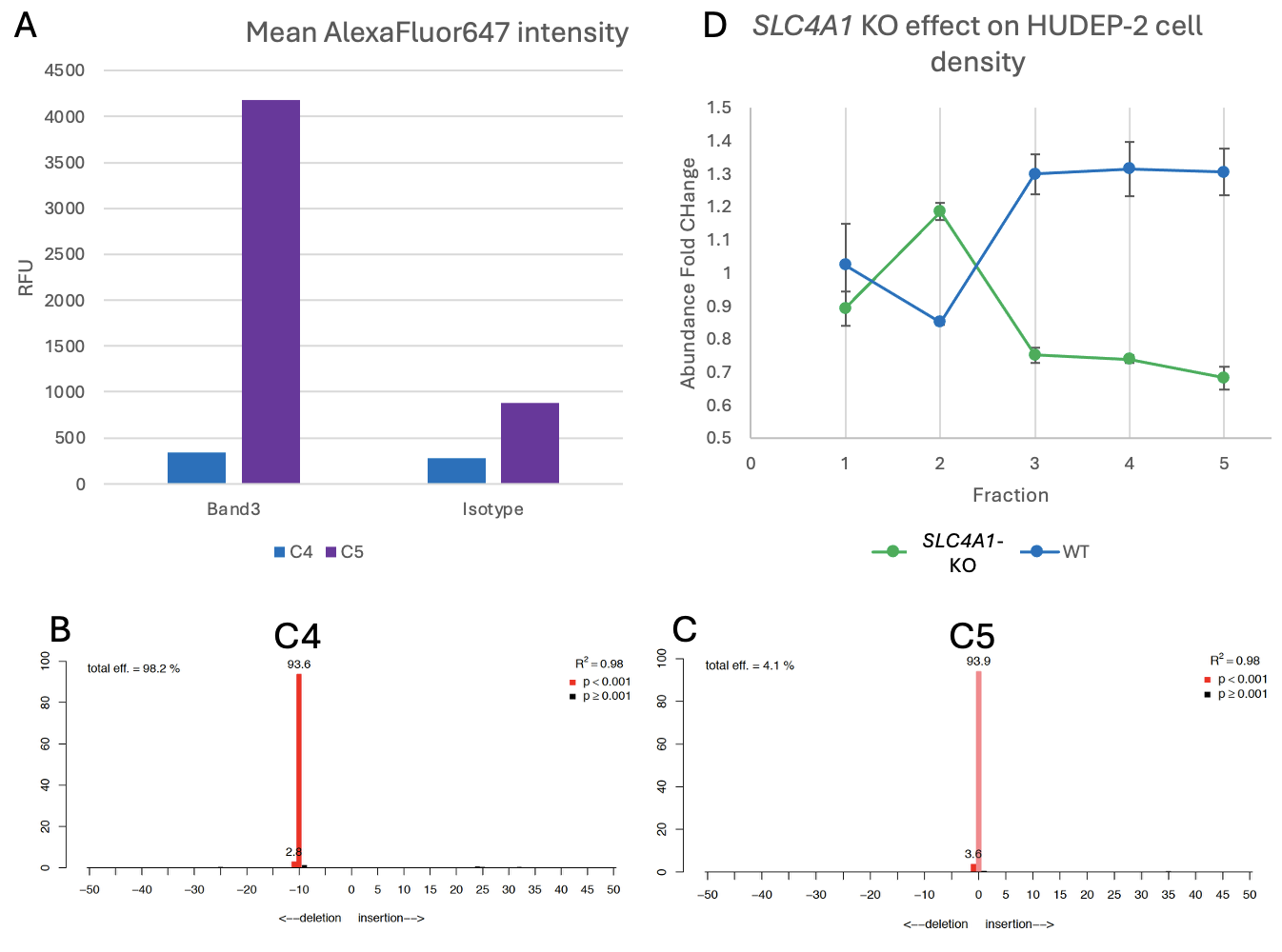

**Supplemental Figure 2. Effect of Band3 (*SLC4A1*) deletion on HUDEP-2 cell density.** (**A**) Immunofluorescence analysis of Band3-*SLC4A1* expression in two different clones, C4 (Band3-*SLC4A1*-KO) and C5 (WT). (**B-C**) TIDE analysis (see **Methods**) of the region targeted by the gRNA used to introduce edits in the *SLC4A1* gene. (**D**) Proportion of WT (C5) and Band3-*SLC4A1*-KO cells (C4). We stained Band3-*SLC4A1*-KO cells with CellTracker Green and WT cells with CellTracker Blue. We mixed the cells and applied them to the Percoll gradient. We assessed the proportion of green and blue cells in each gradient fraction by flow cytometry. The proportion of Band3-*SLC4A1*-KO cells decreases in the denser fractions (from left-to-right).

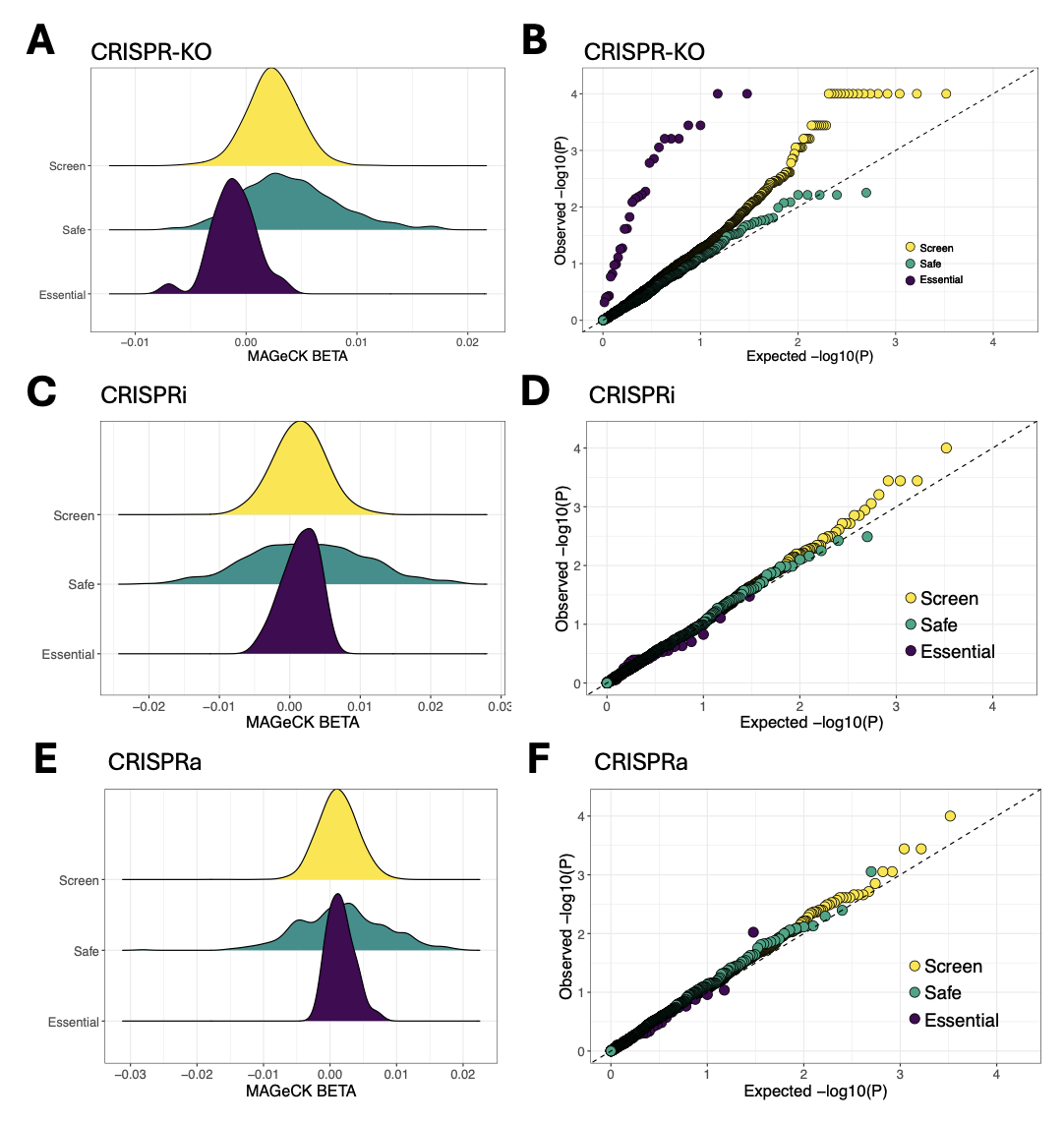

**Supplementary Figure 3. Proliferation results of the HUDEP-2 density screens using three Cas9 modalities and a library of 17,219 gRNA.** Density distributions of effect sizes (BETA, estimated using MAGeCK) for the (**A**) CRISPR-KO, (**C**) CRISPRi and (**E**) CRISPRa screens. The corresponding quantile-quantile (QQ) plots are in **B**, **D** and **F**. In **A**, the BETAs for the gRNA that target essential genes are shifted towards the left compared to the rest of the gRNA, indicating their depletion in the screen. The test statistics for the gRNA that target essential genes is also inflated in **B**. We do not see the same effect in **C-F** because we did not include gRNA that target the promoter of essential genes in our library.

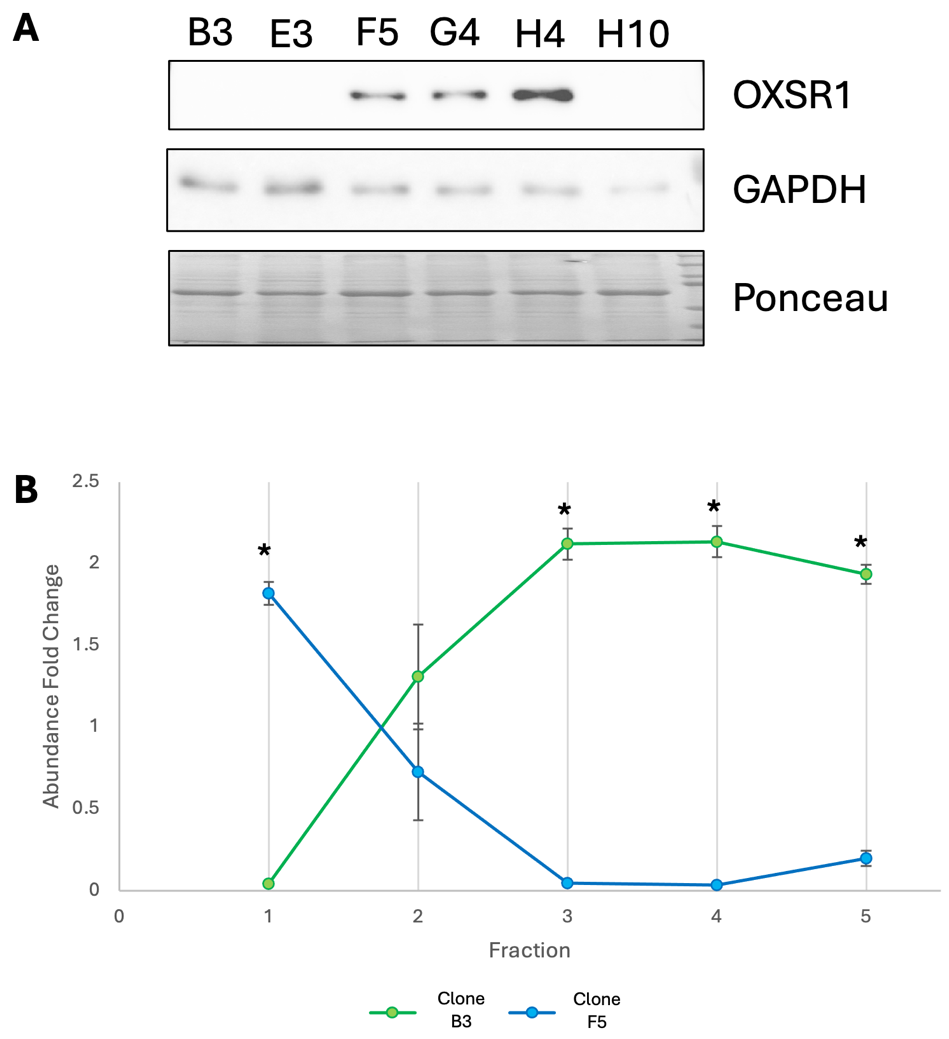

**Supplementary Figure 4. HUDEP-2 clones that do not express *OXSR1* have increased density.** (**A**) Immunoblotting of protein samples from six different HUDEP-2 clones derived from the population of cells that received the *OXSR1* targeting gRNA. (**B**) Proportion of clone B3 and clone F5 cells in each Percoll gradient fractions. We stained B3 cells with CellTracker Green and F5 cells with CellTracker Blue. We assessed the proportion of green cells and blue cells in each fraction by flow cytometry. Cells without OXSR1 (B3, green) have increased density (i.e. more abundant in lower fractions of the Percoll gradient). *P-value <0.05.

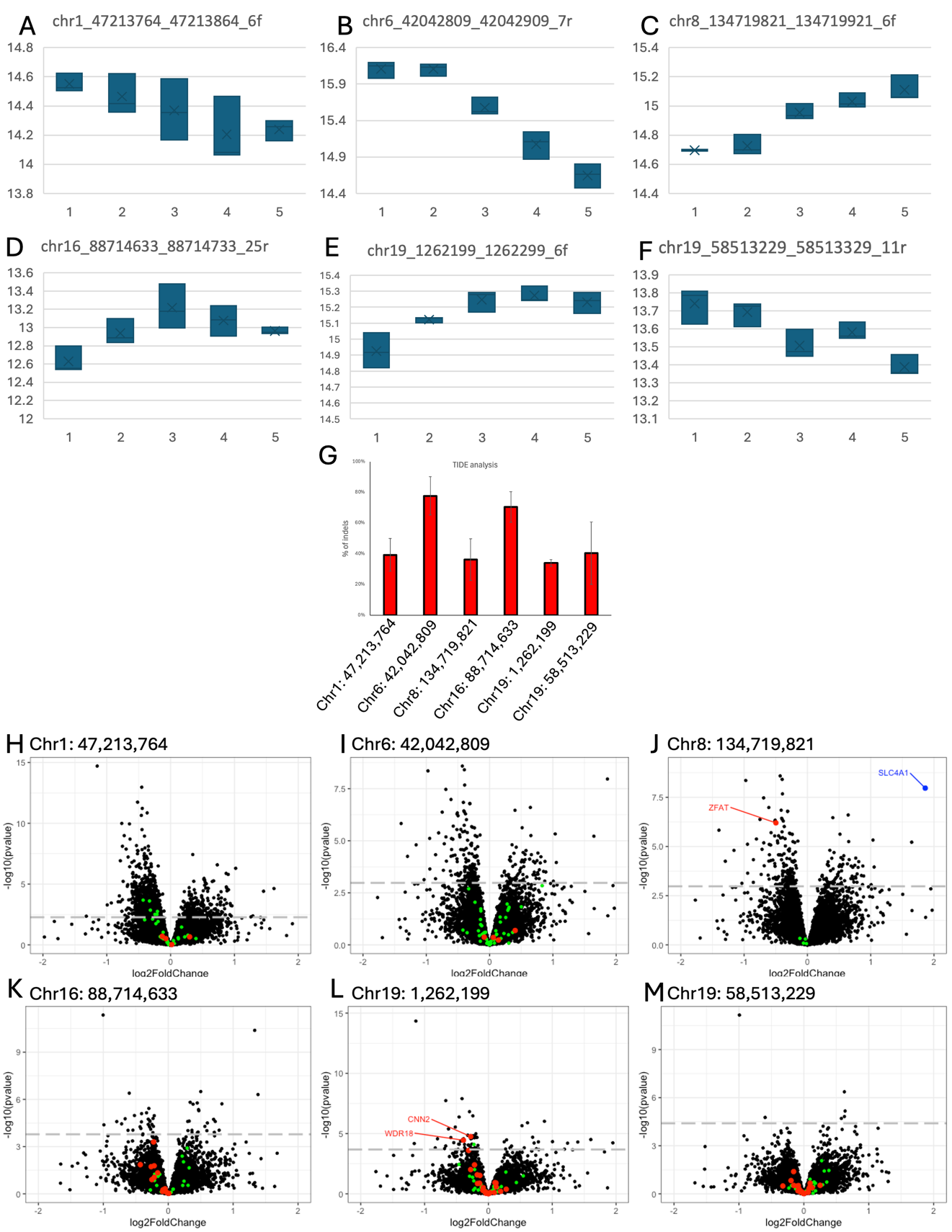

**Supplementary Figure 5. Transcriptomic characterization of six GWAS variant-targeting gRNA that impact the density of HUDEP-2 cells in the Percoll gradient screens.** (**A-F**) Results from the validation screen for the six gRNA that we selected for RNA-sequencing. Three gRNA decrease density (**A**, **B**, **F**) and three increase density (**C**, **D**, **E**). x-axis, Percoll gradient fractions; y-axis, relative gRNA abundance. (**G**) Percentage of indels detected by TIDE analysis in the HUDEP-2 cell populations produced for the RNA-sequencing experiments. (**H**-**M**) Volcano plots showing differentially expressed genes in HUDEP-2 cell populations that received a gRNA targeting regions in **A**-**F**. In red and green are genes located in 1-Mb or 5-Mb windows around the target gRNA, respectively. For chr1:47213764 (**H**), significant genes located within the 5-Mb window (green) are: *UROD*, *UQCRH*, *TOE1*, *PRDX1*, *KIF2C*, *MMACHC*. For chr19:1262199 (**L**), the significant gene located within the 5-Mb window (green) is: *BSG*. chr1:47213764-47213864 (rs4926726, *TAL1*); chr6:42042809-42042909 (rs141929558, *CCND3*); chr8:134719821-134719921 (rs13255015, *ZFAT*); chr16:88714663-88714763 (rs142662688, *CTU2*); chr19:1262199-1262299 (rs10414920, *MIDN*); chr19:58513229-58513329 (rs56225452, *SLC27A5*).
